# Micro-biogeography greatly matters for competition: Continuous chaotic bioprinting of spatially-controlled bacterial microcosms

**DOI:** 10.1101/2020.07.12.199307

**Authors:** Carlos Fernando Ceballos-González, Edna Johana Bolívar-Monsalve, Diego Alonso Quevedo-Moreno, Li Lu Lam-Aguilar, Karen Ixchel Borrayo-Montaño, Juan Felipe Yee-de León, Yu Shrike Zhang, Mario Moisés Alvarez, Grissel Trujillo-de Santiago

## Abstract

Cells do not work alone but instead function as collaborative micro-societies. The spatial distribution of different bacterial strains (micro-biogeography) in a shared volumetric space, and their degree of intimacy, greatly influences their societal behavior. Current microbiological techniques are commonly focused on the culture of well-mixed bacterial communities and fail to reproduce the micro-biogeography of polybacterial societies.

Here, fine-scale bacterial microcosms are bioprinted using chaotic flows induced by a printhead containing a static mixer. This straightforward approach (*i*.*e*., continuous chaotic bioprinting) enables the fabrication of hydrogel constructs with intercalated layers of bacterial strains. These multi-layered constructs are used to analyze how the spatial distributions of bacteria affect their social behavior. Bacteria within these biological microsystems engage in either cooperation or competition, depending on the degree of shared interface. Remarkably, the extent of inhibition in predator-prey scenarios increases when bacteria are in greater intimacy. Furthermore, two *Escherichia coli* strains exhibit competitive behavior in well-mixed microenvironments, whereas stable coexistence prevails for longer times in spatially structured communities. Finally, the simultaneous extrusion of four inks is demonstrated, enabling the creation of higher complexity scenarios.

Thus, chaotic bioprinting will contribute to the development of a greater complexity of polybacterial microsystems, tissue-microbiota models, and biomanufactured materials.

## Introduction

Cells do not work alone, but instead function in highly dynamic societies in which members collaborate and/or compete. For example, cells in human tissues are spatially organized, and this patterning has a significant effect on their functionality. Remarkable examples of this are the highly complex multilayered systems found in the kidney, brain, liver, and cancerous tissues.^[1–5]^ A growing body of evidence suggests that the spatial distribution of microbial societies also matters.^[6,7]^

Micro-biogeography refers to the spatial patterns of microbial communities through time and space.^[8,9]^ The emergence of any particular arrangement greatly depends on gradients in the local microenvironment (i.e., variations in temperature, oxygen concentration, pH, and nutrients).^[10,11]^ Importantly, bacteria outside the community also contribute to the generation of these gradients.^[12–14]^

In nature, micro-biogeographies can be as diverse as well as beautiful. For instance, the distribution and composition of human microbiota vary across different body habitats.^[15]^ Factors inherent to specific sites, such as the salivary flow, may also play critical roles in structuring microbial communities across space.^[16]^ For instance, in caries and periodontal pockets, mosaic architectures of biofilms emerge due to the presence of anaerobes in the interior and aero-tolerant taxa on the exterior, creating hedgehog, corncob, and cauliflower microstructures.^[17,18]^ Therefore, an improved understanding of microbiota organization on teeth, for example, may help in developing more efficient dental therapies.^[19]^

The micro-biogeography in the gut is also very complex and dynamic. For example, studies of gnotobiotic animal models suggest that the intestinal microbiota is distributed along the proximal colon due to microscale mixing, challenging the expected occurrence of spatially segregated communities.^[20]^ However, individual health status may influence this spatial accommodation of microbes. For instance, patients suffering from irritable bowel disease may exhibit an increased bacterial concentration on the mucosal surface compared to healthy controls.^[21]^ The prevention of infections and cancer in other mucosal microenvironments, such as the vagina, has also been associated with the maintenance of a protective shell composed of non-pathogenic bacteria.^[22,23]^

In the plant kingdom, trees also host micro-communities with structured micro-biogeographies, such as lung lichens made of bacteria, algae, and fungi. Beautiful associations of algae and bacteria have been observed in lichen cross sections, forming 30 µm wide interspersed lamellae.^[24]^

In biofilms, bacteria form aggregates made of mono- or poly-bacterial species that play distinct roles according to their phenotypes.^[25,26]^ When bacteria at the periphery cause a depletion of available substrates at the interior, the inner cells starve and interrupt the synthesis of metabolites that are vital for their counterparts on the outside. This dynamic leads to spatiotemporal variations in the bacterial community.^[27]^ A location-dependent metabolism has been observed in clonal colonies of *Escherichia coli (E. coli)*, which form subpopulations specialized in either the Krebs cycle or glycolysis according to their spatial position in the community.^[28]^ These metabolic heterogeneities (i.e., the presence of microscale gradients of nutrients and stressors at the local microenvironment) have been partially associated with the expression of different sets of genes.^[23,29]^ Nevertheless, conventional microbiological culture techniques fail to generate these complex microarchitectures of bacteria and substrates, thereby limiting the study of the effects of spatial variations on the societal dynamics of microbial communities.

One strategy to address this issue is to use biofabrication techniques, such as bioprinting and microfluidics-based manufacturing, among others, in microbiology. For example, Hynes et al.^[30]^ accommodated spatially distinct aggregates of *E. coli* and *Salmonella enterica* using a casting-based method and suggested that interactions in this consortium may be influenced by spatial scales. Similarly, Chen et al.^[31]^ used photolithography to create patterns in adhesion polymers at a resolution of 10 µm. The patterns were subsequently used for specific anchoring of *E. coli* at those locations, and the authors then monitored bacterial crosstalk using a reporter gene activated by the high cell concentration in neighboring fronts. Qian et al.^[32]^ used an extrusion-based system to print diverse 3D-geometries with a 200 µm resolution for biomanufacturing purposes. In particular, lattice-shaped scaffolds containing *Saccharomyces cerevisiae* were capable of a continuous synthesis of ethanol not possible when these organisms were cultured in solid layers. This difference presumably arose because the porosity of the latticed scaffolds facilitated mass transfer.

In this contribution, we present a series of proof-of-concept scenarios to show that: 1) the use of chaotic flows induced by static mixers (i.e., continuous chaotic bioprinting^[33]^) is a versatile tool for creating living microsystems with a printing resolution of a few tens of micrometers, and 2) the micro-biogeography of bacterial communities defines their competition or cooperation outcomes. We first demonstrate that inhibition is strongly influenced by the degree of intimacy between two distinct bacterial strains. We then show that even cells from the same species may exhibit either cooperation or competition, depending on their spatial distribution. Finally, we advance this bioprinting approach for creating scenarios where four different bacterial strains can be incorporated into the same construct or where spatial isolation between bacterial consortia can be established, thereby paving the way for further studies in either fundamental or applied science.

## Results

### Characterization of spatial distribution

Chaotic advection, defined as the continuous stretching and folding of materials that yield a chaotic flow, is extensively used in industrial mixing when turbulence is either unfeasible or inappropriate.^[34–36]^ Chaotic advection can be induced by, for example, a Kenics static mixer (KSM)^[36]^, which is an arrangement of motionless helicoidal mixing elements (**Figure 1**A) fixed in a cylindrical housing.

**Figure 1.**
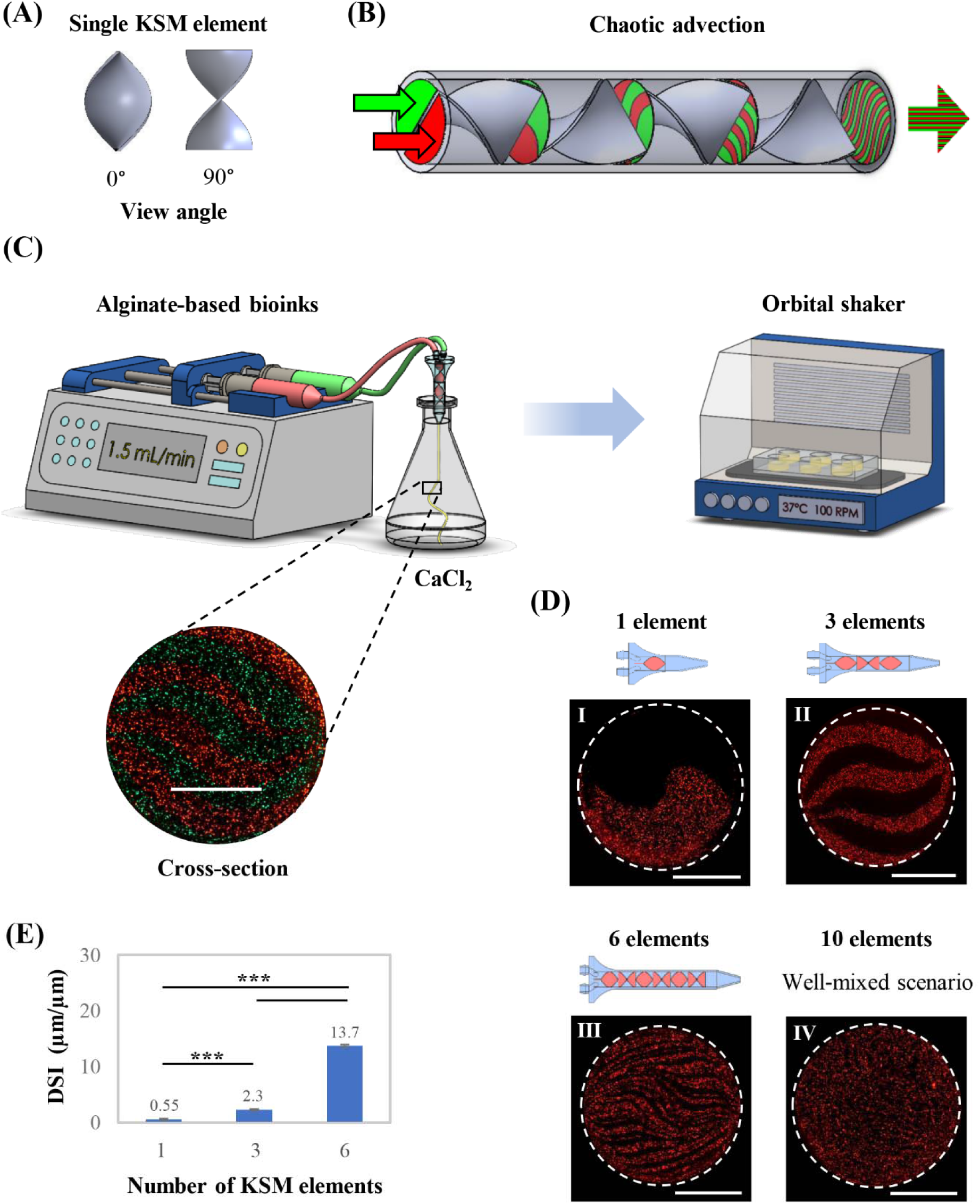
Continuous chaotic bioprinting of fine-scale micro-biogeographies. (A) A single KSM element seen from two view angles. (B) Illustration of a multi-lamellar pattern developed by the successive splitting and folding of bioinks at each mixing element. (C) Schematic diagram of the procedure for bioprinting and culture microcosms-containing fibers using a printhead equipped with 3 KSM elements. The outlet of the KSM printhead must be immersed in the calcium chloride solution. A cross-section of a printed fiber is shown. Scale bar: 500 µm. (D) Printheads containing different numbers of KSM elements and representative micrographs of the micro-biogeographies produced. (E) Quantification of the degree of shared interface (DSI) at each micro-biogeography.**\*\*\****p-value* < 0.001 (n=3).

Recently, our group reported the first use of a KSM for bioprinting spatially organized bacterial communities or mammalian cells at both high throughput (>1 m.min^-1^) and high resolution (∼10 µm).^[33]^ Extrusion of two bioinks through a printhead containing a KSM increases the number of interfaces between them exponentially according to the number of mixing elements (Figure 1B). Interestingly, the printing resolution of our technique, in terms of the thickness of the internal lamellae, can be simply tuned by changing the number of KSM elements in the printhead. In addition, the number of lamellae can be calculated according to the model *s* = 2^n^, where *s* is the number of lamellae and *n* is the number of KSM elements.

Here, we use this biofabrication approach, which we have coined as “continuous chaotic bioprinting,” to provide a precise accommodation of bacterial communities in fine-scale multi-lamellar structures. These bioprinted constructs were then used to assess the impact of the degree of intimacy between bacterial micro-clusters on their social behavior. Our simple printing setup consisted of a KSM printhead, the bioinks (2% alginate containing bacteria), a syringe pump, and a 2% CaCl_2_ bath (Figure 1C). All bioprinting experiments were performed aseptically inside a laminar flow cabinet, and the bacteria-laden constructs were cultivated in suitable growth media at 37°C while shaking at 100 rpm. Our experimental system enables the high-throughput fabrication of fiber-shaped scaffolds 1 mm diameter and containing striations as large as 500 µm or as small as 7 µm.

We first characterized the spatial distribution of our bacterial microcosms. To do this, we co-extruded a suspension of red fluorescent bacteria in pristine alginate ink (2%) with a non-fluorescent pristine alginate ink (2%). Overall, we chaotically printed four different micro-biogeographies: printheads equipped with 1-, 3-, 6-, or 10-KSM elements produced constructs containing 2, 8, and 64 defined lamellae and a homogenous microcosm, respectively (Figure 1D). In principle, the 10-KSM element printhead would render 1024 lamellae of 0.97 μm-thickness, accommodated in a fiber of 1 mm diameter. In this particular setup, the size of the bacteria (approximately 2 μm), which was larger than the average size of the striations, prevented the generation of a layered microstructure.^[37]^

We then established the degree of shared interface (DSI) as a quantitative descriptor of intimacy between bacterial microcolonies (Figure 1E). Here, an interface was defined as the frontier between two striations. The rationale behind the DSI arises from the fact that the inter-material interface is exponentially incremented by chaotic advection.^[33][38]^ The DSI was expressed as the ratio of the total length between lamellae and the fiber perimeter. Figure 1E shows that printheads equipped with 1-, 3-, or 6-KSM elements generate a DSI of 0.55, 2.3, or 13.7 µm/µm, respectively.

Subsequently, we analyzed the reproducibility of the lamellar microstructure along a printed fiber (**Figure 2**A). Figure 2B shows cross-sectional cuts from the same scaffold at different distances and confirms that the striation pattern is highly conserved throughout the whole fiber. Interestingly, like a mirror projection, each red lamella (Figure 2C I) had a practically identical black counterpart (Figure 2C II) in the same cross-section. This phenomenon can be explained in terms of the self-similar nature of chaotic flows, i.e., the repetitive iteration on the same flow manifold throughout *n* number of stretching and folding cycles.^[38][37]^ This self-similarity between lamellae was also evident in simulation results obtained using computational fluid dynamics (CFD) strategies to solve the Navier-Stoke equations of fluid motion (Figure 2C III).^[36]^

**Figure 2.**
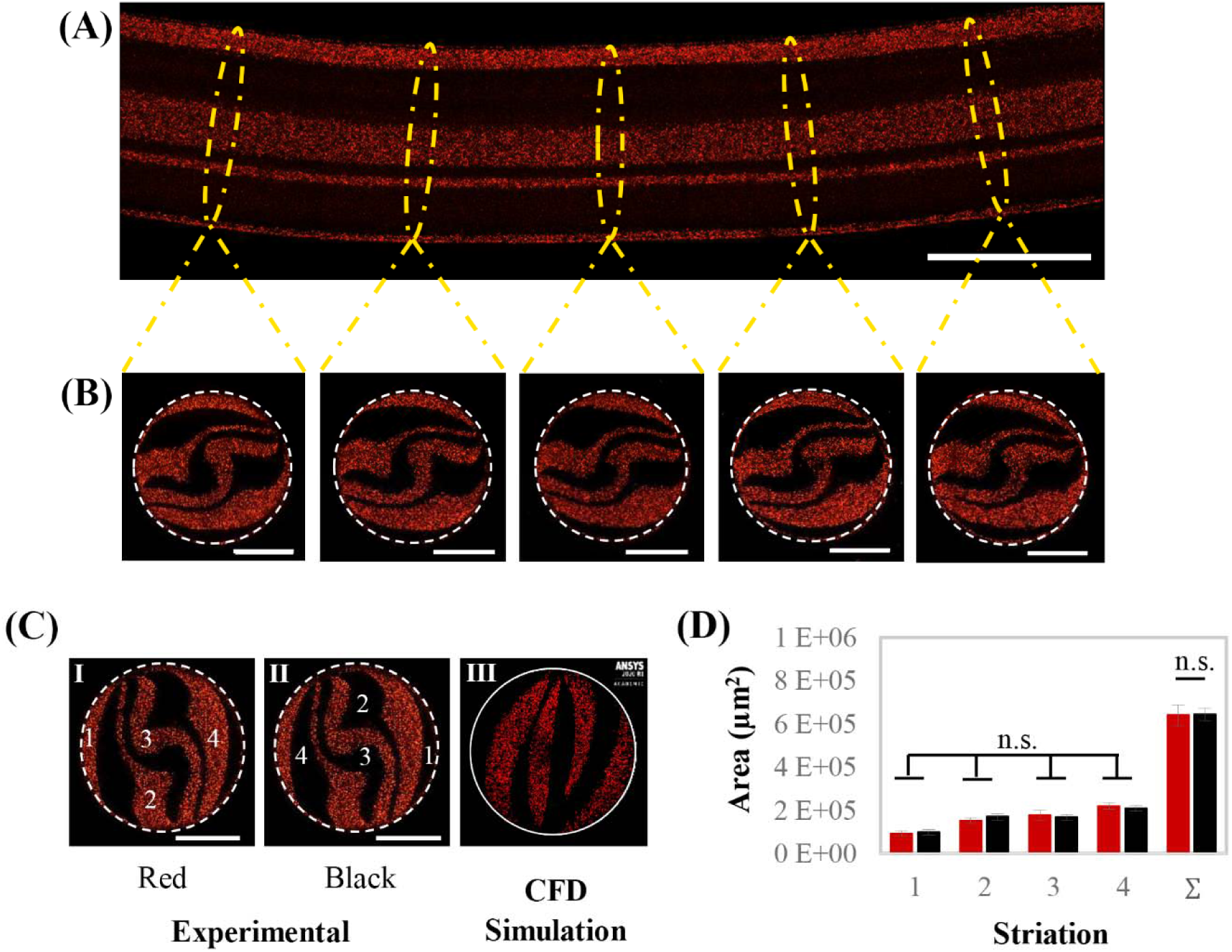
Reproducibility of the lamellar microstructure. (A) Axial view of a fiber printed using 3 KSM elements. Scale bar: 1000 µm. (B) Cross-sectional cuts of the same fiber at different lengths exhibit a conserved multi-lamellar pattern. Scale bar: 500 µm. (C) Mirror-like projections of homologous lamellae marked by the same number (I and II), and CFD simulation of the cross-sectional microstructure after 3 KSM elements (III). (D) The individual and total (∑) areas of red and black striations among 7 cross-sectional cuts obtained experimentally. Non-significant difference (n.s.) at *p-value* < 0.05 (n=7).

We then analyzed a series of cross sections along the fiber and calculated the individual area of homologous lamellae and the cumulative area of the black and red lamellae (Figure2D). We found that the area of the analogous lamellae, (i.e., the symmetric black and red lamellae in the same cross section) is practically equivalent. The areas of analogous lamellae at different cross sections were also equivalent (variance coefficient from 6 to 14%; Supplementary Table 1). The projected cumulative area of the black and red regions at each cross section was also practically identical. This implies that both inks occupy the same amount of territory (surface) in the scaffold; therefore, our bacterial strains would be equally distributed when contained in a chaotically printed micro-biogeography.

### *E. coli* versus *Lactobacillus rhamnosus* GG

In nature, single bacteria form three-dimensional communities, and this spatial conformation influences gene expression and mass transfer of signaling molecules.^[29,39,40]^ Communication and/or synchronized behavior in bacterial communities are conceived as phenomena that mainly depend on the cell density and diversity of the species within the community.^[12,31]^ However, a growing body of evidence suggests a paramount role as well for spatial positioning in bacterial societal behavior.^[6–8,23,41,42]^

In our first biological scenario, a recombinant *E. coli* strain, engineered to produce a red fluorescent protein (EcRFP), and a non-recombinant *L. rhamnosus* LGG (LGG) were used for modeling predator-prey competition. LGG is a probiotic though to suppress overgrowth of pathogenic gut bacteria through diverse mechanisms including interference with pathogen adhesion, secretion of antibacterial compounds (i.e, lactic acid, antibacterial peptides, etc.), and stimulation of the host immune response.^[43,44]^

Prior to the bioprinting process, both bacteria were cultivated for 24 h in fresh medium to achieve a maximum cell density. Each bacterial culture was then centrifuged and placed into a different reservoir containing a mixture of sodium alginate and culture medium. We ensured the reproducibility of our experiments by adjusting the initial cell density of our bioinks to an optical density (OD) at 600 nm of 0.1 and 0.025 for EcRFP and LGG, respectively. These ODs rendered nearly the same number of colony-forming units (CFUs) for each strain just after bioprinting (approximately 7.8 × 10^7^ CFU/mL).

We printed micro-biogeographies using a printhead equipped with 1-, 3-, or 6-KSM elements. These bacterial constructs were cultivated over a 12 h duration at 100 rpm and 37°C in a mixture of Luria-Bertani (LB) and de Man, Rogosa and Sharpe (MRS) media (2:1, v/v), which had been determined as suitable for the co-culture in preliminary experiments. At least three independent experimental runs were performed to evaluate each micro-biogeography.

In all the scenarios analyzed, fluorescence signals in the micrographs decreased steadily over time (**Figure 3**A). **Figure S1** depicts the fluorescence intensity of EcRFP in these bacterial microcosms. We further investigated this trend by assessing the viability of both EcRFP and LGG every 4 h by enumerating CFUs using the agar-plate method. Interestingly, the growth dynamics of both EcRFP and LGG were influenced by DSI (Figure 3B).

**Figure 3.**
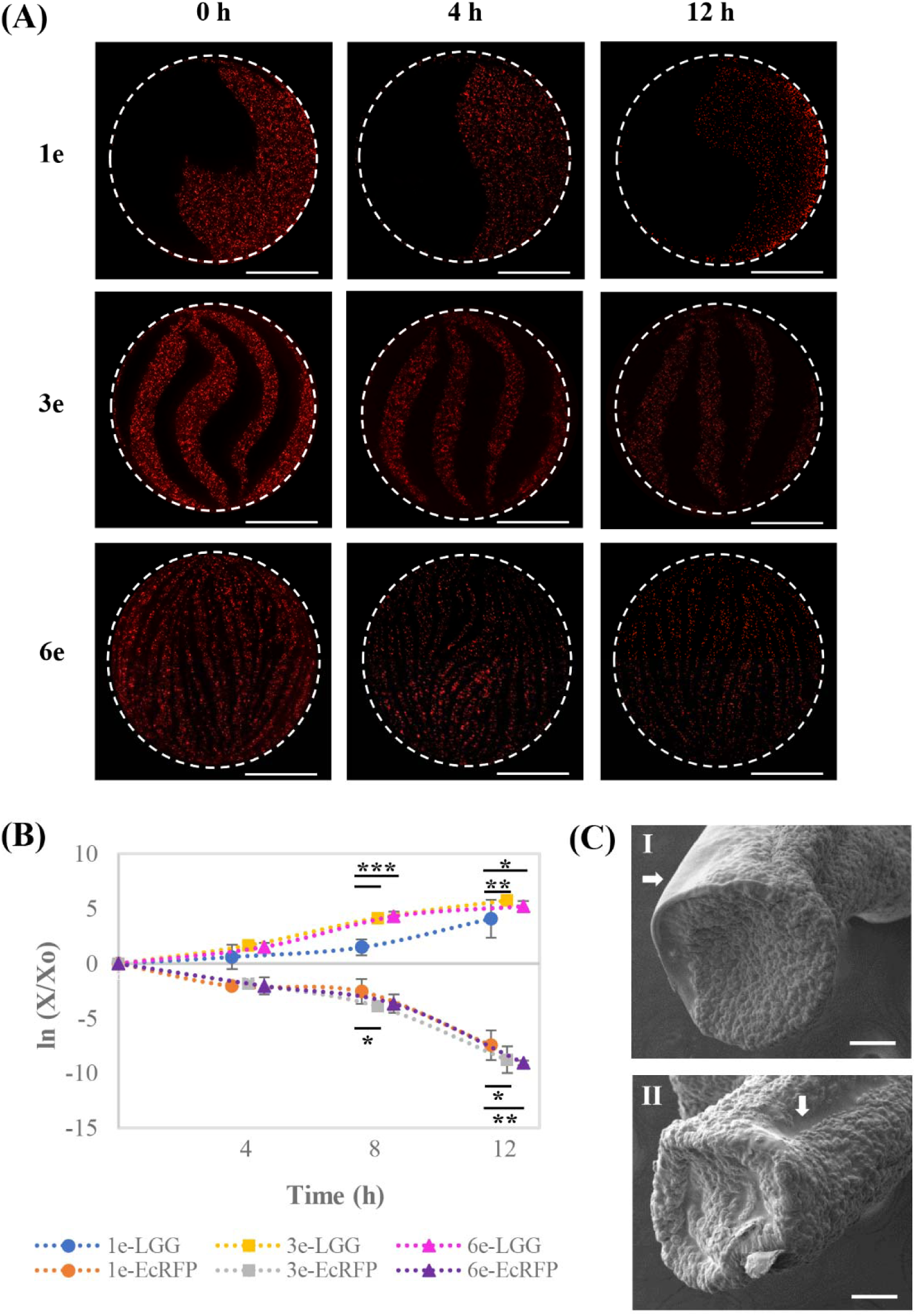
Dynamics of the inhibition of *Escherichia coli* by *Lactobacillus rhamnosus* LGG. (A) Cross-section of the micro-biogeographies containing EcRFP-expressing *E. coli* (red) and LGG (black), chaotically printed using 1, 3, or 6 KSM elements (Scale bar: 500 µm). (B) Co-culture viability over a 12 h duration, normalized by the number of colony-forming units (CFUs) just after bioprinting. (C) SEM micrographs showing the invasion of EcRFP-expressing neighborhoods by LGG, in constructs printed using 1 (I) and 3 (II) KSM elements. **\****p-value* < 0.05; **\*\****p-value* < 0.001; **\*\*\****p-value* < 0.001 (n=6).

The EcRFP viability decreased by the same proportion in all the printed microcosms during the first 4 h of cultivation. However, after 12 h, the inhibition of EcRFP by LGG was less severe in the 1-KSM-fabricated constructs (DSI=0.55 µm/µm) than in the 6-KSM-fabricated microcosms (DSI=13.7 µm/µm; *p-value* < 0.01). This suggests that the lowest DSI was more favorable than the higher ones for EcRFP culture. We hypothesize that specific competition mechanisms of LGG, which were inactive or inefficient at the lowest DSI, were probably triggered at higher degrees of intimacy with EcRFP, thereby inducing a stronger inhibition of this prey. In fact, the proximity of competitors has been suggested to regulate toxin secretion or inhibitors in bacterial communities^[25]^

Concomitantly, LGG experienced a stunted growth during the first 8 h when co-cultured in 0.55 µm/µm DSI micro-biogeographies (fabricated using a 1-KSM printhead) when compared to growth in microcosms printed with 3- and 6-KSM printheads (*p-value* < 0.001). In addition, this tendency was also noticed at 12 h (*p-value* < 0.05; *p-value* < 0.01). The lactate produced by LGG is also noxious itself^[45]^. High local concentration of this metabolite may have contributed to a slowing down of the proliferation of LGG since this lactate-producing strain was confined in a single lamella.

Bacteria can use both contact-dependent and independent competition mechanisms.^[46–48]^ In the first case, a bacterium secretes toxic substances directly into the cytoplasm of a member of a different species; contact is therefore mandatory and DSI is directly relevant. In the second case, bacteria secrete specialized metabolites that diffuse into the microenvironment, where they interfere with the metabolism of susceptible microbial individuals. Distance is also relevant here since diffusion is inversely proportional to the square root of distance.^[33]^ DSI is also highly relevant, since diffusive processes occur more effectively across structures with high perimeter-to-area ratios. Chemical and physical gradients at the local microenvironment play important roles in the dynamics of mixed bacterial communities, so they might influence gene expression and, consequently, growth dynamics.^[27,49]^ In addition, spatial distribution has been suggested as a key driver of gene expression due to the micro-scale concentration differences at distinct locations within a microbial consortium.^[27,29,40]^ Furthermore, spatial segmentation may mitigate the proliferation of specific bacterial species due to changes in local concentrations of signaling molecules.^[23,50]^

Micro-biogeographies printed using a 1-KSM printhead provide only one frontier for competition, allowing a safer establishment of microcolonies, far away from the battlefront. Consequently, the survival outcome in this microcosm may be mainly influenced by the lethality of contact-independent weapons.

Another interesting observation was that the fluorescence intensity in the 1-KSM-fabricated microcosms decreased near the shared border after 12 h (Figure 3A). In fact, Figure 3C I shows that much of the EcRFP neighborhood was invaded by LGG. This trend was also noticeable in the micro-biogeographies printed using a 3-KSM printhead (Figure 3C II). However, when looking at the surface of the fibers (indicated with white arrows), a relatively mild invasion was detected in comparison to the deeper regions. This phenomenon might reflect an effect of the higher oxygen concentration at the construct surface since the proliferation of *E. coli* is potentiated by aerobic microenvironments.

Recently, Song et al.^[51]^ assessed the capability of microencapsulated LGG to either disrupt or inhibit biofilms formed by *E. coli*. Exponential reduction of the biofilm was observed as early as 4 h of co-culture. In addition, the authors found that the 3D-microenvironment stimulated the release of inhibitory molecules by LGG, thereby reducing the transcriptional activation of the *luxS* quorum-sensing pathway in *E. coli*. Analogous reports using the same prey (*E. coli*), but a different predator (i.e., *Bdellovibrio bacteriovorus*), have suggested that this bacterial species exhibits an enhanced persistence when its microcolonies are placed far away from its enemy and, in particular, at the periphery of the micro-landscape.^[52]^

In conclusion, this predator-prey scenario demonstrates that the inhibition dynamics was greatly dependent on the DSI between EcRFP and LGG. Importantly, a high number of viable LGG cells has been suggested to represent a determining factor in the effectiveness of medical interventions using this probiotic strain.^[53]^ Therefore, studies that use LGG to suppress the growth of other microorganisms (i.e., those related to the activity of LGG against pathogenic bacteria) should consider that micro-biogeography may play paramount roles both in the inhibition dynamics of the prey and in the proliferation of LGG.

### *E. coli* versus *E. coli*

Quantum paradoxes are those in which phenomena at the macroscale differ from those at the atomic scale. Likewise, studies on bacterial communities sometimes reveal astonishing information about dynamics related to micro-biogeography.

In our second proof-of-concept scenario, the same *E. coli* strain used above (EcRFP) was cultivated with an *E. coli* strain producing a green fluorescent protein (EcGFP). At first sight and considering the similarity between the weapons of these two bacterial armies (both share practically the same genetic load), we would not expect any fierce competition between them. However, factors such as limited space and resources may lead to competition among bacterial strains even within the same species.^[25,26,54]^ In this set of experiments, we additionally printed a micro-biogeography using a 10-KSM element to obtain a well-mixed microcosm, as stated before. Prior to the bioprinting process, both EcRFP and EcGFP were cultivated in fresh medium for 24 h. Pellets of the bacteria where suspended in sodium alginate, and the initial cell density of each bioink was adjusted to an optical density of 0.1, measured at 600 nm. The bacteria-laden constructs were cultivated for 48 h in LB medium at 100 rpm and 37 °C. In all of the printed micro-biogeographies, EcRFP and EcGFP exhibited increased fluorescence intensity from 0 to 12 or 24 h, and a subsequent decrease after 48 h of cultivation (**Figure 4**A). **Figure S2** depicts the changes in fluorescence intensities for EcRFP and EcGFP over time.

**Figure 4.**
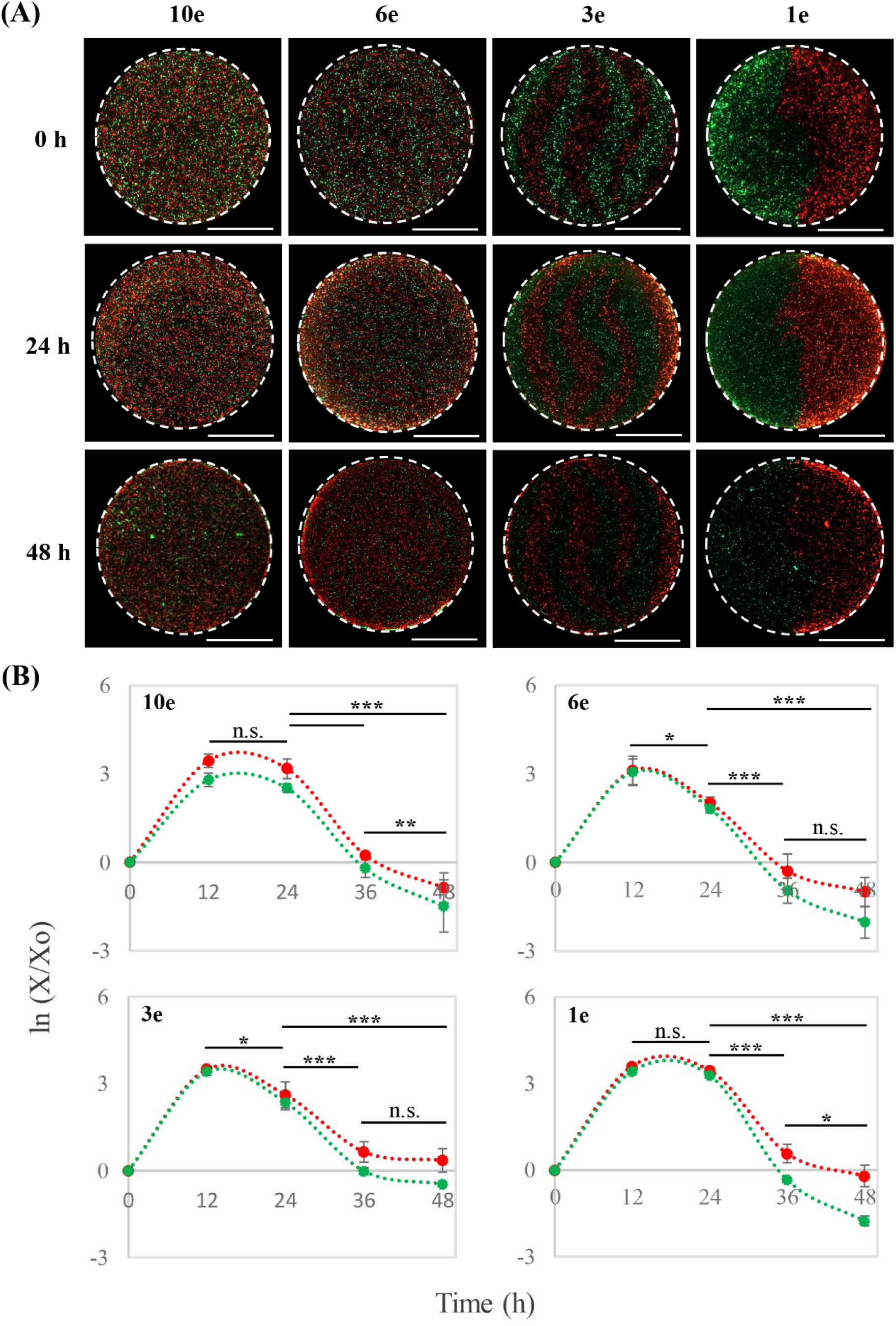
Growth dynamics of the *E. coli* consortia. (A) Cross-section of micro-biogeographies containing EcRFP (red) and EcGFP (green), chaotically bioprinted using 10, 6, 3, or 1 KSM elements (Scale bar: 500 µm). (B) Viability of EcRFP and EcGFP for 48 h, normalized by the number of CFUs just after bioprinting (X_0_). One plot per micro-biogeography. **\****p-value* < 0.05;**\*\****p-value* < 0.001; **\*\*\****p-value* < 0.001 (n=6).

Interestingly, the quantification of the total number of CFUs, using the agar-plate method, revealed that the cooperation dynamics in the community was strongly influenced by the degree of intimacy between its members. We observed a pronounced prevalence of EcRFP over EcGFP throughout cultivation in the 10-KSM-mediated microcosm, i.e., the well-mixed micro-biogeography, (*p-value* < 0.01; *p-value* < 0.001). *In silico* models have suggested that well-mixed microenvironments demand more complex interactions between *E. coli* strains than do segregated systems for supporting co-existence^[55]^, since individual bacteria sense genetic relatedness in their surrounding counterparts in order to establish or avoid cooperation.^[13,14]^ Therefore, a mutualistic behavior is less likely to occur in well-mixed microcosms because the bacteria can freely screen their vicinal homologues, thereby avoiding the risk of being exploited by freeloaders.^[14,56]^

The capability of using available carbon sources and electron acceptors is crucial for determining who dominates who in *E. coli* consortia.^[57]^ Experimental data have also shown that harmonious co-existence between *E. coli* strains is disrupted in well-mixed microcosms.^[58]^ Indeed, one strategy used to promote cooperation in a microbial consortium is to control the spatial position of the microorganisms.^[7,13,41,58]^ However, spatial patterning may not be suitable in specific cases, for example, when inter-strain communication is required for efficient biosynthesis^[42]^, auxotrophy^[59]^, or gene transfer^[60]^.

We then investigated the effect of the spatial accommodation patterns in our *E. coli*–*E. coli* consortium in promoting a sustained growth of both strains. We first analyzed the micro-biogeography in constructs chaotically printed using a 6-KSM element printhead (Figure 4A). In this microcosm, the DSI between EcRFP and EcGFP was 13.7 µm/µm (Figure 1E). Both strains grew at equal proportions. In contrast with the well-mixed condition, the emergence of a dominant strain was not observed (Figure 4B). Both, EcRFP and EcGFP reached a peak of growth at 12 h, with an average magnitude similar to the highest viability of EcRFP in the well-mixed micro-biogeography (3.1±0.1 and 3.4±0.2 in logarithmic scale, respectively). Subsequently, both strains exhibited a death phase starting after 24 h of cultivation.

In a similar fashion, when the microcosm was chaotically printed using 3-KSM elements, both EcRFP and EcGFP grew equitably and steadily for 12 h of cultivation. Nevertheless, the average peak of growth was higher in the consortium printed using 3-KSM elements than using 6-KSM elements (3.5±0.1 vs 3.1±0.1, respectively). This trend was also noticeable at 24 h in constructs printed using either 3- or 6-KSM elements (2.5±0.2 and 1.9±0.2, respectively). Therefore, a smaller DSI (2.3 µm/µm) was more favorable for *E. coli* strains than a larger DSI (13.7 µm/µm) in terms of growth dynamics. Interestingly, in the micro-biogeography printed using 1-KSM element (0.55 µm/µm DSI), neither EcRFP nor EcGFP exhibited a death phase at 24 h. Instead, a continued stationary growth phase was evident (Figure 4B). Our results are consistent with several recent reports that indicate a higher stability in partially segregated communities. For example, the creation of a single shared interface enabled the culture of two *E. coli* consortia for a longer time than when homogenously mixed microenvironments were used.^[61]^ A balanced “chasing” takes place when *E. coli* strains are spatially segregated, thereby boosting ecosystem biodiversity for a longer period.^[58]^

Our results suggest that segregated coexistence, even between bacterial strains from the same species, may facilitate efficient cooperation. The presence of one or more shared interfaces between EcRFP and EcGFP clearly bypassed competition. Remarkably, the microcosm with the lowest DSI evidently facilitated the emergence of a much longer stationary phase for the segregated coculture.

When the same *E. coli* strain is accommodated in spatially segregated subgroups, intra-strain genetic relatedness may diminish over time, forcing each subgroup to consume nutrients at distinct rates and proportions.^[14,62]^ Interestingly, previous reports have shown that microcolonies of *E. coli* may exhibit different growth rates, even when they belong to the same clone, because they adopt distinct metabolic tasks according to microscale gradients of nutrients and metabolites.^[29,40]^ Furthermore, i*n silico* models of *E. coli* have suggested that cells at the edge of a bacterial patch are in charge of expanding the colony boundaries, whereas cells at the interior play different roles, such as cross-feeding.^[49]^ In our system, the degree of competition among our segregated societies appears to be lower in segregated *E. coli* societies than in completely mixed microcosms. As more interfaces exist between the segregated regions, the long-term stability of the community is compromised. While societies fabricated using a printhead with 6-KSM elements reached their peak population in 12 hours and then declined (*p-value* < 0.05), societies that shared one order of magnitude less interface reached a significantly higher population peak during the same time and remained a stable society for 12 additional hours.

The short-term stability of societies printed using 3 and 6 elements might be related to changes in gene expression, and their associated influence in the growth behavior of the community.^[27]^ For example, we hypothesize that additional energy will be invested to maintain the boundaries between both *E. coli* strains (i.e., the red and the green armies) in a segregated microcosm with a higher DSI values. Note also that the rate of growth is significantly lower in the microcosms fabricated with 6 KSM elements than in the ones printed using 3 elements.

Alternative interpretations of our results are certainly plausible. For example, previous reports have suggested that contact-dependent inhibition systems are crucial for cooperation in consortia of either *E. coli* or *Vibrio cholera*. In this ecosystem, each strain naturally adopts the spatial distribution that best fits the community needs.^[26,56,63]^ Our results suggest that, in our *E. coli–E. coli* microcosms, the metabolic necessities at the global level were better addressed at the lowest DSI. Nevertheless, general assumptions should be avoided, since each microbial consortium develops a unique network of metabolic interactions that dictates the community behavior.^[7,64,65]^

We found that the growth dynamics in our *E. coli* consortium can be finely controlled by simply switching the DSI from 13.7 to 2.3 or to 0.55 µm/µm, thereby facilitating the emergence of a stationary growth phase whenever necessary. This straightforward approach may find powerful applications as well in bioproduction engineering and synthetic biology, as segregated ecosystem diversity might make possible the synthesis of added-value compounds and highly complex materials even without redesigning metabolic pathways.^[25,57,66]^

### Tailor-made deposition of multiple inks

Extrusion-based 3D-bioprinters often implement sequential injection protocols for printing multi-material constructs.^[2,3]^ Although remarkably enabling multi-material bioprinting systems have been developed recently, several challenges are still associated with accurate deposition patterns of bioinks at high resolution and fast speed.^[3,67]^ In this final section, we expand the use of our bioprinting system to multi-material printing scenarios that consider the use of more than two inks and enable the facile development of highly complex microbial communities composed of up to four species.

Here, we redesigned our KSM printhead for simultaneous extrusion of four inks from the same nozzle in a simple, continuous, and symmetrical fashion. We achieved this by accommodating four inlets, instead of two, in the SolidWorks design of our KSM printhead. **Video S1** shows that this KSM printhead has a divisor wall that connects the inlet port to the first KSM element. This cap creates two compartments before the first KSM element, each one receiving the feed flow from two inlets. Since our printing technique uses chaotic flows to mix in the laminar regime^[33,36,38]^, all four inks are accommodated in the form of multilayered microstructures with astonishing alienation accuracy (**Figure 5**). The limiting factor in printing resolution will be particle size.^[33]^

**Figure 5.**
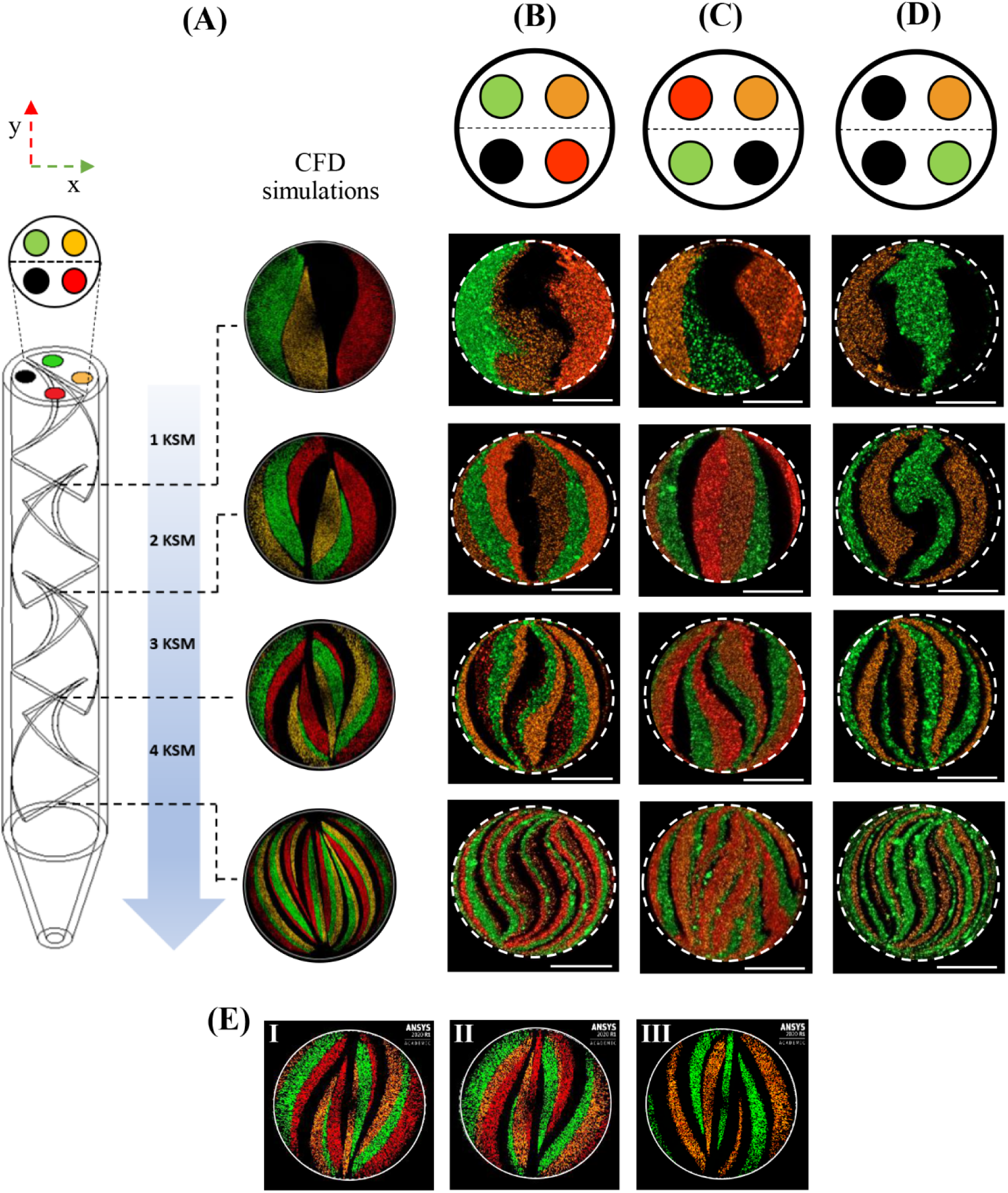
Continuous chaotic printing of multiple inks. (A) Computational simulations showing the evolution of the lamellar micro-structure due to the successive splitting and reorientation of four inks in the KSM printhead. Spatial distribution patterns of alginate inks when (B) the reservoirs of black and green inks were connected to adjacent inlets located at different compartments (divisor wall represented by the dotted line), (C) the black and green inks enter the KSM printhead through inlets positioned at the same compartment, and (D) two black inks were connected to adjacent inletports located at distinct compartments to establish spatial isolation of the green and orange lamellae. (E) Computational simulations of the lamellar patterns obtained when using 3 KSM elements and the connection configurations detailed in B, C, and D.

Our computational simulations show that, when using this four-stream system, the number of internal lamellae in a chaotically printed scaffold increases exponentially according to the model *S*_4*i*_ = 2^*n* + 1^, where *s* is the number of lamellae and *n* is the number of KSM elements in the printhead (Figure 5A). **Video S2** shows CFD simulations of the extrusion process in our multi-port chaotic printing system.

Indeed, our experimental results corroborate the development of the same number of internal lamellae dictated by this model (Figure 5B-D). We found that the spatial organization of the inks in a printed construct could be easily tuned by switching the spatial positioning of each ink at the inlet port. To explain this, consider a XY plane at the top of a KSM printhead (Figure 5), where the X axis is parallel to the divisor wall (represented by a dotted line). When the reservoirs of two inks (2% alginate containing black or green microparticles) are connected to adjacent ports from different compartments (Figure 5B), the inks will not come into contact, regardless of the number of lamellae in the printed construct.

This remarkable obedience of the inks is due to: 1) the architecture and sequential arrangement of the KSM elements, and 2) the laminarity of the flow. Figure 5A shows that adjacent KSM elements have opposite twist directions. Therefore, when the black and green inks are co-extruded across each KSM element in our 4-inlet printhead, they will be split and reoriented to a different compartment, thereby avoiding the emergence of a shared interface between the black and green lamellae. By contrast, when the reservoirs of black and green inks are connected to the inlet ports detailed in Figure 5C, the black and green lamellae will share an interface in the printed fiber because both inks enter the same compartment in the printhead. Consequently, the emergence or absence of shared interfaces between materials, or bioinks, can be precisely defined only by switching the connection order of the reservoirs at the inlet port. These multi-material scaffolds may enable, for example, the fabrication of spatially structured multi-microbial consortia with the presence or absence of shared interfaces.^[24,58,68]^

Finally, implementing the rationale behind Figure 5B, we were able to establish spatial separation between two inks (orange and green) by connecting the reservoirs of two black inks to the adjacent inlets from different compartments (Figure 5D). In this case, the orange and green lamellae exhibited an interspersed repetition in the chaotically printed fiber without any shared interface. The thickness of these physical barriers also decreased due to the incremented number of lamellae. Thus, this type of scaffold may be used to study, for example, the maximum physical separation that allows a syntrophic or auxotrophic consortium to survive^[25,30,42,68,69]^, or to analyze the role of spatial segmentation in horizontal gene transfer.^[59,60,70]^ In addition, our four-stream system would facilitate the accommodation of physical barriers between microorganisms with a micrometer resolution, which might be useful to drive microbial cooperation in biomanufacturing applications. ^[54,64,71–73]^

## Concluding remarks

In this study, we have demonstrated the feasibility of using continuous chaotic bioprinting to fabricate micro-biogeographies with an unprecedented resolution among extrusion-based approaches. To our knowledge, this is the first report to investigate the growth dynamics of microcosms composed of two microorganisms at a wide range of spatial scales, where the DSI is a key factor that defines competition or cooperation outcomes. We postulate that DSI is a suitable quantitative descriptor of the degree of intimacy between members of a bacterial community. Consideration of a complementary parameter related to the spatial position of microcolonies (possibly the average distancing) is also recommended.

In further support of our statement regarding DSI, consider the shared interface between a pair of striations as a battlefront, where members of both armies fight and die to maintain the defined boundaries of their territories. Meanwhile, their counterparts fulfill other duties far away from that location. Consequently, when the area occupied by a specific population of bacteria decreases, we can intuitively expect that a dramatic change in metabolic tasks, such as cross-feeding, might occur. In fact, the feasibility of this analogy has been partly confirmed in previous reports.^[29,40,49]^ Thus, continuous chaotic bioprinting may be an enabling platform in microbial ecology and related scientific fields that will allow the creation of spatially controlled living microsystems at high printing resolution. We envision the utility of chaotic bacterial bioprinting for the development of customized microcosms in the analysis, for example, of the effects of micro-biogeography on the microbial transcriptome or gene transfer. We anticipate that this might have tremendous relevance to the design of probiotic interventions and the understanding of the interplay between different species in complex ecosystems such as the gut, the oral cavity, or the skin. Implications of spatial segregation to phenomena such as antibiotic resistance^[29,74]^ may be rigorously studied in chaotically bioprinted systems. Advanced metabolomic techniques, such as 13C metabolic flux analysis^[28]^, might also be implemented to identify phenomena occurring locally (i.e., within lamellae boundaries).

Nature is a magical and prolific factory of scenarios where spatial distribution matters. The unprecedented development of a four-stream KSM printhead expands the horizons of continuous chaotic bioprinting to diverse and exciting applications. For instance, multi-port chaotic bioprinting may enable the facile creation of the beautiful lamellar microarchitecture of lung lichens, where bacteria, algae, and fungi co-exist in 30 µm wide interspersed lamellae.^[24]^ Other scenarios include the investigation of horizontal gene transfer according to the spatial segmentation of the microenvironment^[59,60,70]^, or the analysis of the influence of spatial micro-organization on the capacity of pathogens to withstand antimicrobials agents.^[29,74]^

We believe that the implications of our findings will be also highly relevant in scientific fields outside microbiology, where additive manufacturing technologies are used in research. Multi-port chaotic printing may facilitate the synthesis of multi-material structures with tailored local properties^[75,76]^, or the mimicry of compositional gradients found in tissues such as the bone and the skin^[3,77]^, among other applications.

## Experimental section

### Printing set-up

In general, our continuous chaotic printing system consisted of a commercial syringe pump (Fusion 200, Chemyx) loaded with 5 mL sterile syringes, sterile plastic hoses, a disinfected printhead containing a specific number of KSM elements (Figure 1C-D), and a flask containing 2% aqueous calcium chloride (Sigma Aldrich). The KSM printheads were printed on a Form 2 SLA 3D printer (FormLabs) using a standard resin (Clear FLGPCL04, FormLabs). We used the design parameters reported in our previous contribution^[33]^ to establish a printhead outlet diameter of 1 mm. A new version of our KSM printhead was designed using SolidWorks, incorporating four-inlet ports instead of two, for simultaneous extrusion of four inks. In this four-stream system, each pair of inlets connects to one of the two compartments inside the printhead. These compartments originate from the cap that connects the base of the inlet ports to the first KSM element (Video S1).

### Bacterial strains and culture conditions

Bacterial cultures were grown in distinct reservoirs for 24 h at 37°C before printing experiments. *E. coli* strains expressing either RFP or GFP were separately cultured in Luria-Bertani (LB) medium (Sigma Aldrich) containing 1 µL/mL of chloramphenicol to retain the recombinant plasmid. *L. rhamnosus* GG (LGG) (ATCC 53103) was grown in de Man, Rogosa, and Sharpe (MRS) medium (Merck).

### Bioink preparation and printing experiments

Bioinks were prepared following this general protocol: approximately 10 mL of each bacterial culture was centrifuged at 8000 rpm for 10 min. The supernatant was discarded, and the pellet was resuspended in 2% sterile alginate solution (Sigma Aldrich) supplemented with suitable culture medium (i.e., 2% LB broth for EcRFP or EcGFP, and 5.22% MRS broth for LGG).

In *E. coli* versus *L. rhamnosus* GG experiments, the initial cell density of the bioinks, in terms of optical density, was adjusted to 0.025 and 0.1 absorbance units for LGG and EcRFP, respectively. In *E. coli* versus *E. coli* experiments, the cell density of both EcRFP and EcGFP was adjusted to 0.1 absorbance units. Subsequently, each bioink was deposited in a distinct sterile syringe and connected to a KSM printhead. The bioinks were co-extruded at a flow rate of 1.5 mL/min at room temperature while the printhead outlet was immersed in a 2% calcium chloride solution. Bacteria-laden constructs were cultured in 6-well plates containing 3 mL of MRS-LB (1:2) medium for *E. coli* versus *L. rhamnosus* GG micro-biogeographies, or LB medium for *E. coli* versus *E. coli* experiments. In both scenarios, the printed fibers were incubated at 37°C under shaking at 100 rpm.

The number of CFU was assessed by disaggregating and homogenizing 0.1 g of sample in 0.9 mL of phosphate buffered saline (PBS). A 100-µ L aliquot of homogenized sample was sequentially diluted in PBS. Sequential dilutions were seeded by duplicate on Petri dishes containing MRS-agar or LB-agar medium. The collected data was multiplied by the dilution factor, and normalized by the number of CFU in constructs just after bioprinting. The results were depicted in logarithmic scale. At least six independent constructs were analyzed at each time point.

In a final set of experiments, we printed multi-color constructs using our four-inlets KSM printhead and 1 part of fluorescent microparticles (Fluor Green 5404; Fluor Hot Pink 5407; or Fluor Sunburst 5410, Createx Colors) in 9 parts of a 2% pristine alginate solution. Fluorescent microparticles were washed as detailed in our previous report.^[33]^

### Microscopy analyses

The microarchitecture and fluorescence of the chaotically printed fibers was assessed using an Axio Observer.Z1 microscope (Zeiss) equipped with Colibri.2 led illumination and an Apotome.2 system (Zeiss). A stitching algorithm, included in the microscope software (Zen Blue Edition, Zeiss), was used for producing wide-field micrographs.

### Characterization of micro-architecture

The DSI of each micro-biogeography was estimated according to Equation 1. The measurements were performed using Image J software, by Fiji. The results were expressed as the average of three independent micrographs per micro-biogeography (*n=3*).

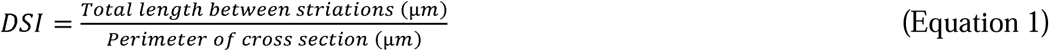

We evaluated the reproducibility of the lamellar microstructure by calculating the area of each black or red striation using Image J software. Once the scale bar was set, each lamella was surrounded using the freehand selection tool. The average of seven cross-sectional cuts(*n=7*) was reported. In addition, the total area of either red or black lamellae in the cross-section was expressed as the sum of individual measurements.

### Scanning electron microscopy (SEM)

SEM images were obtained using a variable pressure scanning electron microscope EVO/MA25 (Zeiss). Briefly, printed fibers were sequentially incubated with 4% formaldehyde and 4% paraformaldehyde for 15 min each, and then washed with PBS. The fibers were then successively dehydrated in an ethanol gradient (i.e., using 25, 50, 75, and 95% ethanol in water) for 1 h. The samples were then coated with gold and visualized at high vacuum mode.

### Computational simulations

Computational fluid dynamics simulations were implemented using ANSYS Fluent 2020 software. The 3D geometries of the systems were discretized using a fine mesh of triangular elements, and a mesh refinement procedure was conducted to ensure convergence of results. Using this mesh, the Navier Stokes equations of motion were solved at each node in laminar flow using a transient state implementation. A fluid density of 1000 kg m^-3^ and a viscosity of 0.1 kg m s^-1^ were used.^[33]^ No-slip boundary conditions were imposed in the fluid flow simulations.

### Statistical analysis

All statistical analyses were performed using GraphPad Prism 8. Biological data were presented as the mean ± SD from at least six bacterial constructs (n=6). Based on two-way analysis of variance (ANOVA) and Tukey multiple comparisons, differences between data were considered statistically significant at **\****p-value* < 0.05, **\*\****p-value* < 0.01, or **\*\*\****p-value* < 0.001.

## Acknowledgments

CFCG and EJBM gratefully acknowledge financial support granted by CONACyT (Consejo Nacional de Ciencia y Tecnología, México) in the form of Graduate Program Scholarships. GTdS acknowledges the funding received from CONACyT, L’Oréal-UNESCO-CONACyT-AMC (National Fellowship for Women in Science, Mexico), and UC-MEXUS. MMA and GTdS acknowledge funding provided from CONACyT. YSZ acknowledges the funding granted by the Brigham Research Institute. This research has been partially funded by the Tecnologico de Monterrey. We gratefully acknowledge the experimental contributions of Felipe López-Pacheco, Norma Alicia Garza-Flores, Carolina Chávez-Madero, Alan Roberto Márquez-Ipiña, and Everardo González-González to this work.

## Author contributions

CFCG, EJBM, MMA, and GTdS designed the study. CFCG and EJBM analyzed the data. CFCG, EJBM, MMA, and GTdS wrote the manuscript. CFCG, EJBM, and KIBM performed all the bacterial bioprinting experiments. DAQM conducted all the computational simulations. CFCG, EJBM, GTdS, and DAQM designed the four-inlet KSM printhead. CFCG, EJBM, and LLLA conducted the printing characterization of the four-stream system. CFCG, EJBM, and DAQM prepared the illustrations. JFYdL fabricated the printheads by stereolithographic 3D printing. GTdS, MMA, and YSZ edited the final versions of the manuscript. All the authors read, commented on, and approved the manuscript.

