## Supporting information for "Micro-biogeography greatly matters for competition: Continuous chaotic bioprinting of spatially-controlled bacterial microcosms"

Centro de Biotecnología-FEMSA

Tecnologico de Monterrey

Monterrey 64849, NL, México

^#^Authors contributed equally.

D.A. Quevedo-Moreno, Prof. G. Trujillo-de Santiago.

Departamento de Ingeniería Mecatrónica y Eléctrica

Escuela de Ingeniería y Ciencias

Tecnologico de Monterrey

Monterrey 64849, NL, México

Prof. Y.S. Zhang

Division of Engineering in Medicine, Department of Medicine

Brigham and Women’s Hospital

Harvard Medical School

Cambridge 02139, MA, USA

Prof. M.M. Alvarez

Departamento de Bioingeniería

Escuela de Ingeniería y Ciencias

Tecnologico de Monterrey

Monterrey 64849, NL, México

J.F. Yee-de León

Delee Corp., Corp.

Mountain View 94041, CA, USA

**Table S1.** Comparison of the surface area of analogous striations within a cross section printed using a 3 KSM printhead.

| **Area per striation (µm^2^)** | | | | | | | | |
| --- | --- | --- | --- | --- | --- | --- | --- | --- |
| **# Lamella** | **Red** | | | | **Black** | | | |
|  |  | **Mean** | **SD** | **CV (%)** |  | **Mean** | **SD** | **CV (%)** |
| **1** |  | 9.22E+04 | 1.25E+04 | 1.36E+01 |  | 9.87E+04 | 1.29E+04 | 1.30E+01 |
| **2** |  | 1.51E+05 | 1.32E+04 | 8.78E+00 |  | 1.70E+05 | 1.48E+04 | 8.69E+00 |
| **3** |  | 1.78E+05 | 2.37E+04 | 1.33E+01 |  | 1.67E+05 | 1.29E+04 | 7.76E+00 |
| **4** |  | 2.19E+05 | 1.35E+04 | 6.16E+00 |  | 2.09E+05 | 1.26E+04 | 6.05E+00 |
| **Total** |  | 6.40E+05 | 4.86E+04 | - |  | 6.44E+05 | 2.88E+04 | - |

*Mean values, standard deviation (SD), and coefficient of variation (CV).*

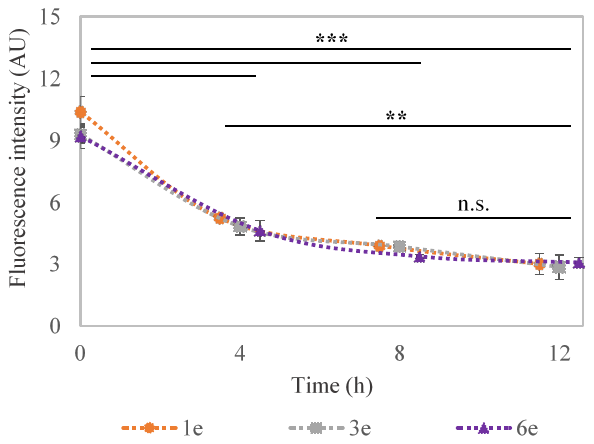

**Figure S1.** Fluorescence intensity (FI) of *E. coli* (EcRFP) in co-culture with *L. rhamnosus* (LGG) for 12 h. These micro-biogeographies were printed using 1, 3 or 6 KSM elements to obtain 2, 8 or 64 internal lamellae. Gray scale values were obtained from micrographs (red channel) to calculate FI, g. FI was expressed as the mean gray value (*n=3*). *******p-value* < 0.01; ********p-value* < 0.001. The decrease in FI was directly proportional to the decrease in viability of EcRFP.

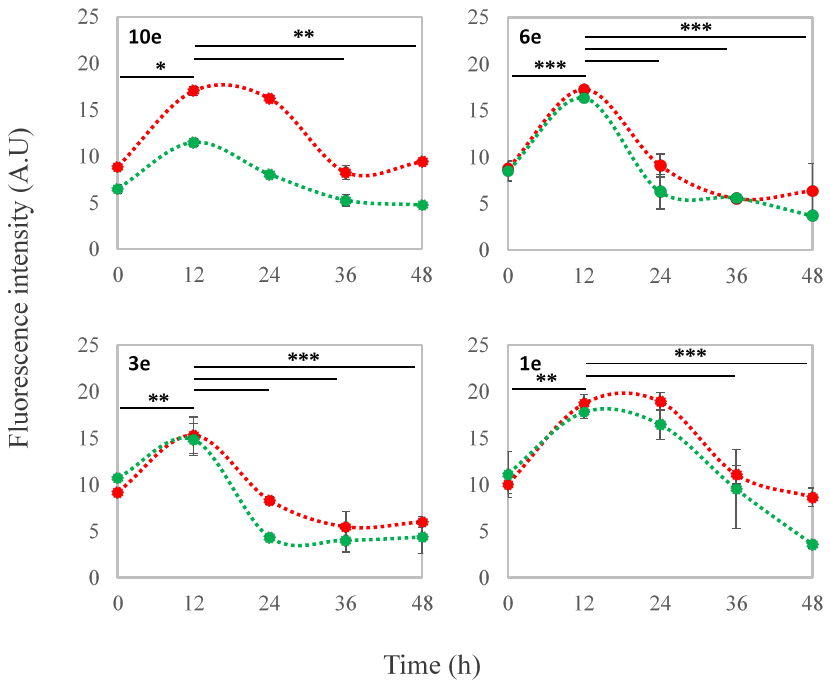

**Figure S2.** Fluorescence intensity (FI) of *E. coli* (EcRFP) and *E. coli* (EcGFP) in co-culture for 48h. These micro-biogeographies were printed using 10, 6, 3 or 1 KSM elements. The printhead containing 10 KSM elements produced a well-mixed micro-biogeography. Gray scale values were obtained from individual micrographs (red or green channels) to calculate FI. FI was expressed as the mean gray value (*n=3*). ******p-value* < 0.05; *******p-value* < 0.01; ********p-value* < 0.001. Fluctuations in FI were closely related to viability of both EcRFP and EcGFP over time.
